# Localizing Unsampled Regions in Partially Sampled Pathogen Phylogenies Using Machine Learning

**DOI:** 10.64898/2026.11.18.767590

**Authors:** Marzieh Khodaei, Nathaniel Hupert, Arindam RoyChoudhury

## Abstract

**Motivation:** Pathogen phylogenies are useful for monitoring the evolution of pathogens and for detecting new variants, as it provides information that can aid public health decisions and outbreak response. Since not all infections are sequenced, certain pathogen lineages may be missing or only poorly represented in reconstructed phylogenies. Identifying these gaps could help to determine where additional data collection is required.

**Results:** We developed a computational method to locate regions of missing samples from the observed tree alone, without sequences from those samples. A machine learning model predicts which branches lead toward the missing region. Beam search then identifies three representative paths and estimates a candidate most recent common ancestor (MRCA) of the missing region for each path. We tested this framework on 5,000 simulated phylogenies where we controlled which tips were deleted. On an independent test set, the best candidate recovered an average of 89.9% of deleted tips, increasing to 90.7% when all three candidates were combined. Predicted and true MRCAs lay on the same ancestral–descendant lineage in 91.3% of cases. We further evaluated the approach on an empirical SARS-CoV-2 phylogeny with introduced missingness. By identifying potentially underrepresented lineages, this framework could help real-time identification of missing infections and improve outbreak response.

**Availability and Implementation:** Code and reproducibility instructions will be made publicly available at https://github.com/TaraKhodaei/phylo-missingness-localization.

**Supplementary Information:** Supplementary data are available online.

## Introduction

Pathogen phylogenies have now become an essential part of genomic surveillance, offering an evolutionary framework for understanding how infectious agents appear, diversify, and spread. Platforms such as Nextstrain have allowed genomic and epidemiological information to be combined so that the evolution of pathogens can be tracked in near real time (Hadfield et al., 2018), and phylogenetic analyses of evolutionary relationships and phylodynamic analyses of population and transmission dynamics played a key role during the COVID-19 pandemic in describing the introduction of the virus, the dynamics of its lineages, its geographic spread, and the emergence of new variants (Attwood et al., 2022; Vöhringer et al., 2021). Yet the information contained in these phylogenies depends on which infections are detected and managed to be sequenced.

Often some infections are missing from the data, and sequencing coverage varies across geographic regions and surveillance systems (Brito et al., 2022). As a result, some pathogen lineages may be only partially represented or entirely missing from reconstructed phylogenies, leading to gaps in the observed evolutionary history. For example, wastewater sequencing has revealed previously undetected SARS-CoV-2 transmission that was not identified at the time (Karthikeyan et al., 2022). These cases show that missing infections can leave pathogen diversity unobserved in reconstructed phylogenies.

Missing data can influence phylogenetic analyses depending on its extent and location within the tree. For instance, if certain regions or host populations are sampled less frequently, their pathogen lineages may be underrepresented or absent from the tree (Lemey et al., 2020; Song et al., 2026). During the COVID-19 pandemic, rural and minority communities were underrepresented in SARS-CoV-2 sequencing data, potentially leaving gaps in reconstructed phylogenies (Kim et al., 2023). In the present study we look at a particular type of missing data in which the missing genomes are found among the descendants of a specific ancestral lineage, leading to missing data that is localized in a phylogenetic tree. The question then is whether the part of the phylogeny in which these missing samples are concentrated can be determined entirely on the basis of the observed tree.

Pathogen phylogenies are most informative when sampled genomes adequately represent the pathogen lineages circulating in the population. For SARS-CoV-2, low testing rates and uneven sampling across locations and time have been shown to delay detection of emerging variants and bias estimates of their prevalence (Han et al., 2023). These limitations are significant because genomic data inform outbreak investigation, transmission control, and strain selection for vaccine development (Armstrong et al., 2019). Additionally, pathogen phylogenies help the evaluation of newly detected lineages for characteristics such as transmissibility, disease severity, immune escape, and impacts on diagnostics or therapeutics, which influence vaccines and other control strategies (Oude Munnink et al., 2021). Lineages that are poorly represented in genomic data may be overlooked. Localizing the concentration of missing samples within the phylogeny enables the framework to identify specific lineages that may require ongoing monitoring or additional investigation. This information could provide useful context when considering future vaccination, testing, or other public health responses.

Locating the region of missingness is challenging because missing genomes provide no sequence information from which to determine their phylogenetic position directly. Previous phylodynamic work has noted this limitation for unsequenced pathogen samples, whose ancestral attachment to the sampled phylogeny is not directly informed by sequence data (Vaughan et al., 2019). Other recent approaches have addressed different aspects of missing information in pathogen phylogenies; for example, ECHO estimates the number of unsampled infections using phylogenetic and epidemiological information (Doig and Colijn, 2026). These developments point to an ongoing challenge: figuring out where missing samples are clustered in the observed phylogeny when there is no sequence data.

Incomplete sampling occurs when only a fraction of the infected hosts are sampled and thus appear in the reconstructed phylogeny. Currently, the various approaches deal with this by incorporating either the unsampled infections or the sampling process itself into the models used for epidemic dynamics, transmission, and phylogeography. Methods based on phylogenies take account of incomplete sampling when estimating epidemic and population dynamics from phylogenies (Stadler, 2009; Vaughan et al., 2019). Transmission-reconstruction methods also accommodate unsampled infections. For example, TransPhylo reconstructs transmission histories from partially sampled outbreaks and can infer unsampled intermediate hosts (Didelot et al., 2017), whereas SCOTTI includes non-sampled hosts within a structured-coalescent framework (De Maio et al., 2016). More recent extensions of TransPhylo incorporate additional epidemiological information to improve transmission inference in partially sampled outbreaks (Carson et al., 2025). The phylogeographic methods overcome the limitations associated with sampling by taking into account the unsampled diversity and the variations in sampling intensity when reconstructing the geographic or host-state histories (Lemey et al., 2020; Song et al., 2026).

Computational phylogenetics has also addressed issues with missing taxa or incomplete sequence data, in addition to methods designed for pathogens. Missing-taxon placement and tree-completion methods infer positions for taxa absent from a tree when information about those taxa comes from observed traits or other phylogenetic trees (Revell et al., 2015; Mai and Mirarab, 2022). Another localization problem is phylogenetic placement. In this case, methods like EPA-ng, UShER, and EPIK are used to find where a query sequence fits within a reference phylogeny (Barbera et al., 2019; Turakhia et al., 2021; Romashchenko et al., 2023). Such methods depend on sequence information from the query taxon, but this information is not available when the missing descendants have not been observed at all. Methods for predicting missing phylogenomic sequences deal with a different type of missingness, namely the absence of genes or partitions for taxa which are still represented elsewhere in the dataset (Darriba et al., 2016). Current methods either include missing observations in larger evolutionary or epidemiological models, or they use information linked to the missing taxa themselves. Determining the ancestral location of missing data by making use of the structural features of the observed phylogeny is still a separate inference problem.

Machine learning and structured search are increasingly used in phylogenetic inference. Predictive models help prioritize topology-changing moves during phylogenetic tree search (Azouri et al., 2021). Wang et al. combined deep-learning-based topology scores with a modified stepwise-addition procedure using beam search, a search algorithm that retains only a fixed number of the highest-scoring candidate trees at each step for further exploration (Wang et al., 2023). Recent studies have looked at using learned traversal of phylogenetic trees and reinforcement learning to explore tree neighborhoods (Collienne et al., 2026; Camerota Verdú et al., 2026). These works demonstrate that predictive models can work with structured search to examine large phylogenetic spaces while keeping promising options.

Here, we introduce a computational framework for localizing the region of missingness in rooted pathogen phylogenies. Given an observed phylogeny, the framework uses structural features of the tree to infer candidate paths leading toward the missing region. At each internal node, a supervised classifier estimates the descendant branch most likely to lead toward the missing region. This process is applied recursively as candidate paths are extended through the tree. The resulting predictions guide a beam search that preserves several plausible root-to-node paths instead of selecting a single traversal at each branching decision. From these candidate paths, we calculate beam support based on how often each node appears and use it to select three distinct representative paths. For each representative path, the largest decrease in beam support is used to estimate a candidate most recent common ancestor (MRCA), which helps position the predicted missing region along the path. The framework is evaluated using controlled simulations with known localized tip deletions, which enables direct assessment on an independent test set. Its application is further examined using an empirical SARS-CoV-2 phylogeny under controlled localized deletion. This framework offers an interpretable method for identifying where missing samples are most likely concentrated in relation to an observed pathogen phylogeny.

We provide a new approach for identifying potentially missing pathogen lineages using only the observed phylogenetic tree. This information could be used to rapidly and efficiently deploy additional data collection efforts to populations and/or regions potentially experiencing excess as-yet undetected infections.

## Methods

### Problem Formulation

Let *T*_*F*_ denote the complete rooted phylogenetic tree. To simulate localized incomplete sampling, an internal lineage is selected at random and a subset of its descendant tips is removed, while the remaining descendants are retained. Pruning the resulting tree yields the observed phylogeny, denoted by *T*_*O*_.

Since only part of the selected lineage is removed, the missing sequences remain localized within a single evolutionary region of the observed tree. Let

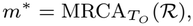

where *ℛ* is the set of retained descendants originating from the selected lineage. The node *m*^*\**^ therefore represents the true evolutionary source of the localized missing region in the observed tree.

The objective is to identify this missing region using only the observed phylogeny. Rather than directly predicting a single node, the proposed framework estimates one or more root-to-node paths that are most consistent with the missing evolutionary signal. The terminal node of each predicted path is interpreted as a candidate source of the missing region.

For simulated data, the deleted tips and the corresponding true root-to-*m*^*\**^ path are known and are used only for training and performance evaluation. During inference, the framework operates solely on the observed phylogenetic tree. The overall workflow of the proposed localization framework is summarized in Fig. 1.

**Figure 1.**
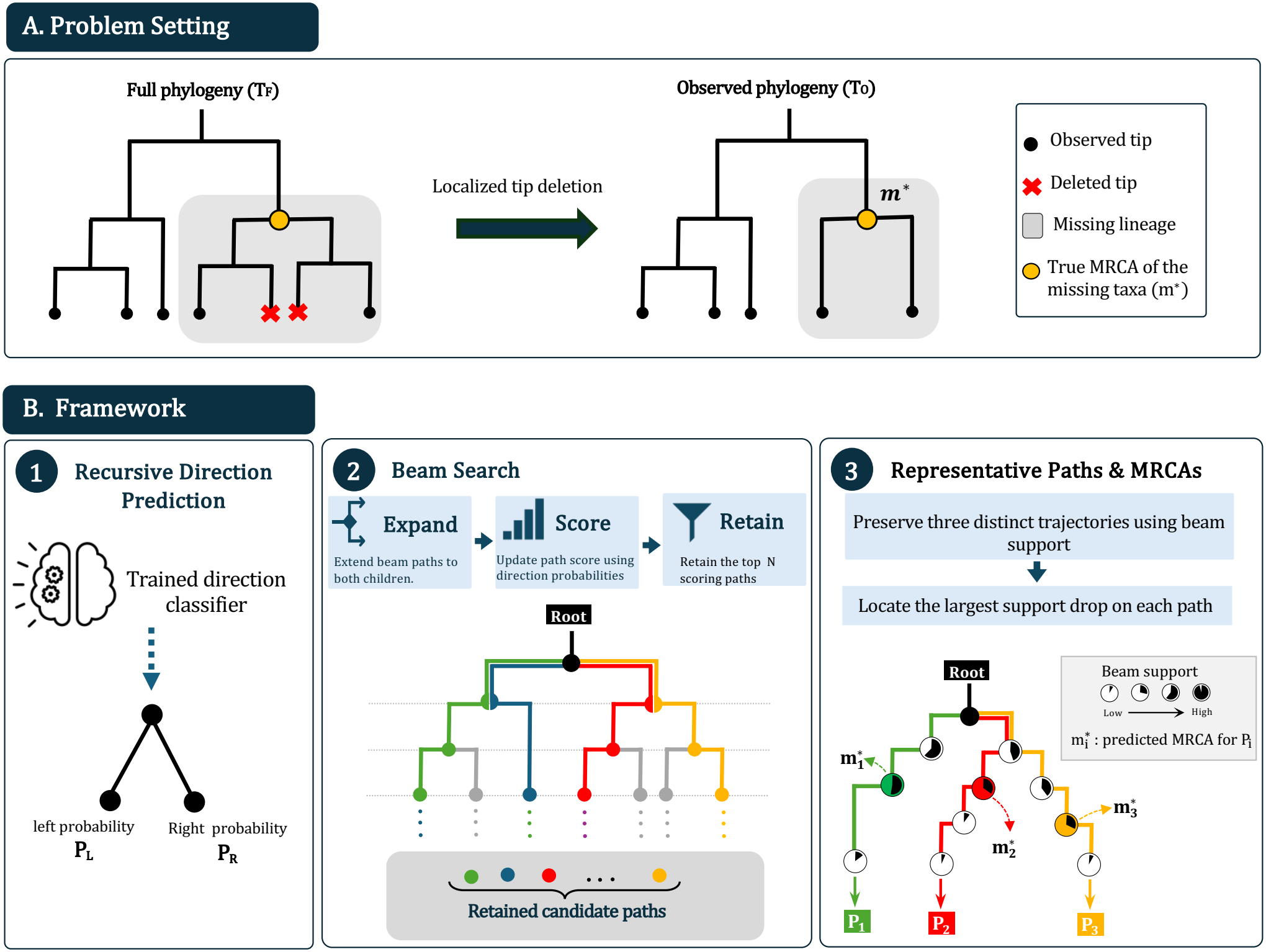
Workflow for localizing phylogenetically clustered missingness. (A) Problem setting. Localized removal of descendant tips from the complete rooted phylogeny, *T*_*F*_, produces the observed phylogeny, *T*_*O*_. The true MRCA of the missing taxa, *m*^*\**^, denotes the ancestral source of the affected lineage. In simulated data, the deleted tips and *m*^*\**^ are known only for model training and performance evaluation, whereas localization is performed from *T*_*O*_ alone. (B1) Recursive direction prediction. At each internal node, the trained direction classifier uses structural characteristics of the two descendant subtrees to estimate *P*_*L*_ and *P*_*R*_, representing the relative evidence that the missing region lies along the left or right direction. (B2) Beam search. The directional probabilities guide recursive expansion of competing root-to-node paths. Candidate paths are updated according to their cumulative scores, and only the highest-scoring paths are retained at each iteration, preserving multiple plausible localization trajectories without exhaustive tree traversal. (B3) Representative paths and MRCAs. Recurrence of nodes across the retained beam paths defines beam support. Beam support is used to select three distinct representative paths, *P*_1_, *P*_2_, and *P*_3_, reducing redundancy while preserving distinct candidate paths to the missing region. Along each path, the largest decrease in beam support identifies a candidate MRCA, 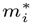, yielding up to three candidate ancestral locations for the missing region.

### Training and Evaluation Dataset Generation

To develop and evaluate the proposed framework, we generated simulated phylogenetic datasets with known missing-region locations. Empirical SARS-CoV-2 phylogenies informed the simulation tree sizes and heights. We reconstructed these phylogenies from complete, high-coverage genomes obtained from GISAID EpiCoV (Khare et al., 2021) for 35 California counties during January 1 to 7, 2021. Instead of using the empirical trees directly for model training or evaluation, we used their tip-count and tree-height distributions to guide the simulated datasets.

Complete rooted phylogenetic trees containing 50–150 tips were generated under four constant-rate birth–death regimes. Tree sizes were sampled using a combination of uniform sampling across the 50–150-tip range and sampling informed by the empirical county-tree sizes. We then rescaled each simulated tree’s branch lengths to match a target height sampled from the empirical tree-height distribution. This approach preserves a realistic evolutionary scale while enabling controlled generation of observed trees for supervised learning and evaluation.

Let *T*_*F*_ denote a simulated complete rooted phylogenetic tree with tip set *ℒ*_*F*_ . For each tree, an eligible internal (non-root) node *u* was randomly selected as the target of localized missingness. The set of descendant tips of *u* is

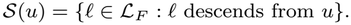

A subset of descendant tips,

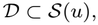

was randomly selected for deletion, while the remaining descendants were retained. The empirical inputs and simulation settings are summarized in Table 1.

**Table 1.**
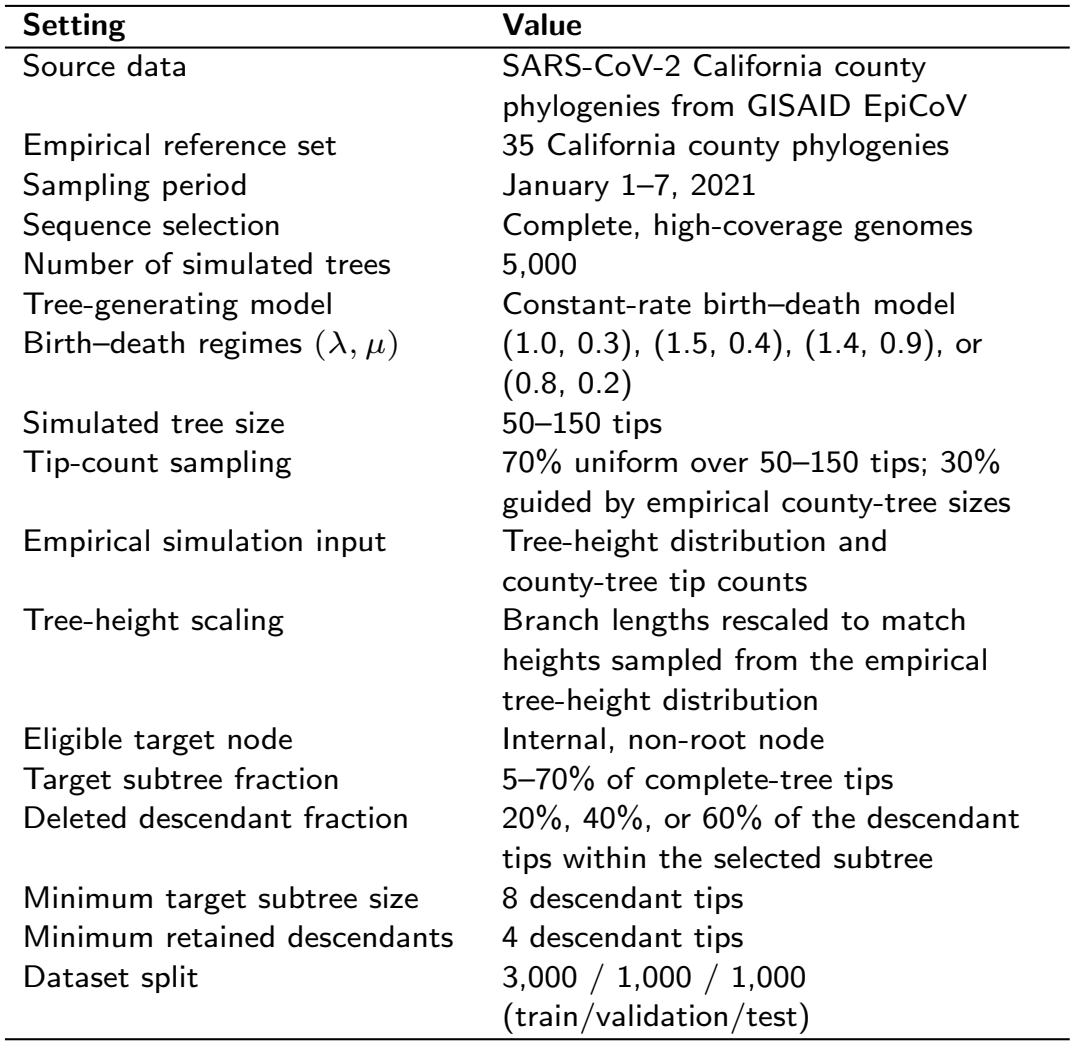
Empirical inputs and simulation settings used to generate the training, validation, and test datasets.

| **Setting** | **Value** |
|---|---|
| Source data | SARS-CoV-2 California county phylogenies from GISAID EpiCoV |
| Empirical reference set | 35 California county phylogenies |
| Sampling period | January 1–7, 2021 |
| Sequence selection | Complete, high-coverage genomes |
| Number of simulated trees | 5,000 |
| Tree-generating model | Constant-rate birth–death model |
| Birth–death regimes (λ, μ) | (1.0, 0.3), (1.5, 0.4), (1.4, 0.9), or (0.8, 0.2) |
| Simulated tree size | 50–150 tips |
| Tip-count sampling | 70% uniform over 50–150 tips; 30% guided by empirical county-tree sizes |
| Empirical simulation input | Tree-height distribution and county-tree tip counts |
| Tree-height scaling | Branch lengths rescaled to match heights sampled from the empirical tree-height distribution |
| Eligible target node | Internal, non-root node |
| Target subtree fraction | 5–70% of complete-tree tips |
| Deleted descendant fraction | 20%, 40%, or 60% of the descendant tips within the selected subtree |
| Minimum target subtree size | 8 descendant tips |
| Minimum retained descendants | 4 descendant tips |
| Dataset split | 3,000 / 1,000 / 1,000 (train/validation/test) |

The observed phylogeny was obtained by pruning the deleted tips,

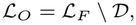

while preserving branch lengths. Because the deleted tips and the selected target lineage were known during simulation, the true MRCA and corresponding root-to-MRCA path could be determined automatically and used for supervised model training and performance evaluation.

Finally, the simulated observed phylogenies were randomly divided into independent training, validation, and test sets for model training, parameter tuning, and unbiased performance evaluation, respectively.

### Recursive Direction Prediction

The localization problem was decomposed into a sequence of binary routing decisions along the observed phylogeny. At each internal node *v*, the objective is to determine which of its two child branches is more likely to continue toward the true localization node *m*^*\**^. The recursive direction model therefore learns the local evolutionary direction required to navigate the tree one branching decision at a time.

Training labels were generated automatically from the simulated datasets. For every internal node on the true root-to-*m*^*\**^ path, the descendant branch containing *m*^*\**^ was assigned label 1, whereas the alternative branch was assigned label 0. Thus, each branching event contributed a pair of supervised training examples.

Each candidate child branch was represented by a 52-dimensional feature vector describing its structural characteristics within the observed phylogeny (Supplementary Table S1). The representation integrated three complementary sources of information: (i) intrinsic properties of the candidate subtree, including its size, branch-length characteristics, and internal balance; (ii) its evolutionary position within the observed tree; and (iii) comparative descriptors characterizing structural asymmetries between the candidate and its sibling branch.

A supervised binary classifier was trained to estimate

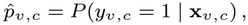

where **x**_*v,c*_ denotes the feature vector associated with candidate child branch *c* at decision node *v*, and

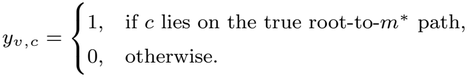

During inference, both child branches are evaluated independently at every decision node. Their predicted probabilities provide local routing scores that guide the beam-search procedure described in the following subsection.

### Beam Path Generation

The recursive direction model provides only local probabilities for branch selection. A traversal that always picks the most probable branch at each step might fail to reach the correct localization path if it makes an early wrong choice. In order to make the procedure more reliable, we use beam search, which simultaneously examines several possible paths through the observed phylogeny.

A beam is a fixed-size set of the highest-scoring candidate root-to-node paths retained during the search. We begin at the root of the observed tree. For each path in the beam, we use the recursive direction model to check which child branches at the end of the path can be extended. Following each such extension, the newly created paths are ranked, and only those with the highest scores within the specified beam width are kept.

Let

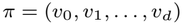

denote a candidate path from the root to node *v*_*d*_. Its score is computed as

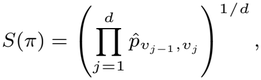

where 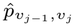 is the recursive direction probability assigned to the selected child at node *v*_*j−*1_. The geometric mean summarizes the average confidence of the routing decisions along the path while avoiding systematic bias against longer paths.

This search continues until no retained path can be expanded or the maximum search depth is reached. In the final implementation, a beam width of 50 produces up to 50 high-confidence candidate localization paths for each observed phylogeny. These beam paths are subsequently aggregated to identify representative localization paths.

### Representative Path Selection

Beam search generates candidate paths with high predicted probabilities of leading toward the missing region. The framework then checks how consistently each node is supported among these paths and makes use of this information to find representative localization paths.

Let

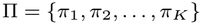

denote the set of retained beam paths. The support of a node *v* is defined as

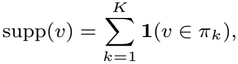

where **1**(*·*) is the indicator function. Therefore, supp(*v*) is the number of beam paths that pass through node *v*.

Node support indicates the number of retained beam paths that go through each node; nodes with high support are used by a large number of possible paths, and a reduction in support shows where these paths split off into different branches. We apply this approach in order to select three representative paths. The first of these follows the branches that have the highest support, and the second and third are taken from the strongest alternative branches, taking into account both their support and the points at which they diverge from the first path. These three paths are used as possible localization trajectories for subsequent MRCA estimation.

### MRCA Estimation

Each representative path shows a possible evolutionary route to the missing region. Let

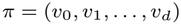

denote a representative path, where each node *v*_*i*_ has beam support supp(*v*_*i*_).

Because the representative paths might overlap close to the root, when estimating the MRCA for each path the process begins at the first node that is not common to the other paths chosen. Within this particular segment of the path, the candidate MRCA is taken to be the node that comes right after the greatest decrease in beam support; if several decreases are equal then the deepest node is chosen. A decrease in beam support indicates that fewer candidate paths continue through that branch, reflecting where the paths diverge. This provides a heuristic for estimating the location of the MRCA.

The same rule is applied separately to each representative path, with one candidate MRCA being selected for each path and this resulting in up to three candidate MRCAs for each observed phylogeny.

### Performance Evaluation

We assessed performance on an independent test set consisting of 1,000 observed phylogenies that had been generated with the simulation settings summarized in Table 1. Initially, the full phylogenies included between 50 and 150 tips, the number of tips varying from one phylogeny to another rather than being split into groups of fixed size. Following the localized deletion, the observed phylogenies had between 34 and 149 tips, with each phylogeny losing 1 to 48 tips (the median number being 5). The test set included 309, 358, and 333 phylogenies in which 20%, 40%, and 60% of the descendant tips in the selected target subtree, respectively, were deleted. Since we knew the deleted tips and their corresponding true MRCA for each of the simulated phylogenies, we were able to assess the localization accuracy directly.

For each observed phylogeny, the proposed framework gives up to three representative localization paths together with their corresponding MRCA estimates. The predictions were assessed according to the proportion of deleted tips that were recovered within the predicted area of missing tips. For each observed phylogeny, the highest recovery rate among the three predictions was taken as the main measure of localization performance.

We considered recovery to be complete if all of the deleted tips had been recovered, partial if only some of them had been recovered, and absent if no deleted tips had been recovered. In order to examine the complementary information offered by the multiple predictions, we also looked at the union of the deleted tips that had been recovered across the three representative paths in the cases of partially recovered phylogenies.

## Results

To examine whether regions of missingness can be located from the structure of a phylogeny alone, we developed and tested the framework using simulated trees with known missing samples. We generated 5,000 rooted phylogenies under different birth–death settings, with tree sizes and heights informed by empirical SARS-CoV-2 data. For each tree, we introduced localized missingness by randomly removing some of the tips descending from a selected internal node, while retaining other descendants of the same lineage. The resulting observed trees were divided into training, validation, and test sets. Using only the observed tree, the framework identifies three candidate paths toward the missing region and estimates a possible most recent common ancestor (MRCA) along each path. The original trees provide the true locations of the missing samples for evaluating these predictions.

We assessed performance on 1,000 independent test phylogenies by measuring how many deleted tips were recovered within the predicted regions and how closely the candidate MRCAs matched the true ancestral locations. We also tested the framework on an empirical SARS-CoV-2 phylogeny with introduced missingness.

### Localization Performance

Table 2 shows how well the proposed framework performed on an independent test set of 1,000 observed phylogenies. For each tree, we first measured recovery using the prediction that found the largest proportion of deleted tips among the three reported localization paths. With this approach, the average recovery across the test set was 89.9%.

**Table 2.** Localization performance on the independent test dataset. Combined recovery was computed from the union of deleted tips recovered by the three representative localization paths.

| **Metric** | **Result** |
|---|---|
| Test phylogenies | 1,000 |
| Complete recovery | 883 (88.3%) |
| Partial recovery | 30 (3.0%) |
| No recovery | 87 (8.7%) |
| Mean recovery using the best prediction | 89.9% |
| Mean recovery using the combined predictions | 90.7% |

We also measured recovery after combining the deleted-tip sets found by all three predictions. This raised the average recovery to 90.7%, which shows that the extra representative paths found deleted tips that the best single prediction missed in some trees.

Complete recovery was obtained for 883 phylogenies (88.3%), partial recovery for 30 phylogenies (3.0%), and no recovery for 87 phylogenies (8.7%).

### Direction Model Performance

The recursive direction model looks at each internal node in the observed phylogeny and predicts which descendant branch is more likely to have the missing lineage. Beam search then uses these predictions to find possible paths to the missing region. Because of this, the direction model’s accuracy is important for overall localization performance.

Model performance was evaluated using 43,556 recursive direction instances extracted from 5,000 simulated phylogenies and partitioned into independent training, validation, and test datasets. Multiple tree-based learning algorithms were evaluated, and the final model was selected exclusively according to its validation performance. Table 3 summarizes the performance of the highest-performing classifiers.

**Table 3.** Performance of the highest-performing recursive direction classifiers. The final model was selected exclusively according to validation accuracy.

| **Learning algorithm** | **Training (%)** | **Validation (%)** | **Test (%)** |
|---|---|---|---|
| HistGradientBoosting | 79.9 | **79.4** | **78.9** |
| Random Forest | 94.2 | 79.0 | 78.5 |
| Extra Trees | 84.5 | 78.4 | 78.4 |

HistGradientBoosting had the highest validation accuracy (79.4%) and performed similarly on the independent test set (78.9%) (Table 3). Random Forest and Extra Trees achieved much higher training accuracies, but this did not lead to better results on the validation or test sets, suggesting that both models were more prone to overfitting. The similar validation and test accuracies of HistGradientBoosting also suggest that its performance was consistent across unseen phylogenies. Based on these results, we selected HistGradientBoosting as the recursive direction model for the subsequent beam-search localization and MRCA estimation analyses.

### Evaluation of Representative Path Selection

We tested the representative path selection strategy on an independent test set to assess how well it could summarize beam-search results and keep different localization hypotheses separate. For each phylogeny, beam search produced 12 to 50 candidate localization paths, with an average of 35.1. We then used these paths to create three representative paths for subsequent MRCA estimation.

Table 4 compares our representative path selection method with simply choosing the three highest-ranked beam paths. The representative paths recovered more of the true missing lineage and had considerably less overlap with one another than the top three beam paths.

**Table 4.** Comparison of representative path selection and raw top-three beam paths. *Lineage preservation* is the mean fraction of the true root-to-MRCA path retained by the selected paths. *Path overlap* is the mean pairwise shared-prefix overlap among the selected paths (lower values indicate less redundancy). *Best recovery* is the mean percentage of deleted tips recovered by the highest-performing path for each phylogeny. *Combined recovery* is the mean percentage of deleted tips recovered after combining the deleted tips recovered by all three selected paths.

| **Metric** | **Raw top 3** | **Representative** |
|---|---|---|
| Lineage preservation | 0.740 | **0.779** |
| Path overlap | 0.293 | **0.136** |
| Best recovery (%) | 82.7 | **89.9** |
| Combined recovery (%) | 82.8 | **90.7** |

The improvements resulted in a great deal better localization performance. Using the proposed representative paths, the framework achieved a recovery of 89.9%, increasing to 90.7% when the deleted tips recovered by all three predictions were combined. In comparison, directly selecting the three highest-ranked beam paths achieved recoveries of 82.7% and 82.8%, respectively.

### MRCA Localization Accuracy

The proposed MRCA estimation strategy was evaluated on the independent test dataset by comparing the predicted and true MRCAs for each observed phylogeny. The predicted MRCA exactly matched the true MRCA in 12.7% of test phylogenies. When we considered the distance between the two nodes, 34.8% of predictions were within one edge of the true MRCA, and 57.1% were within two edges.

The predicted MRCA was located on the same ancestral– descendant lineage as the true MRCA in 91.3% of test phylogenies. As shown in Fig. 2, most predicted MRCAs were located at ancestral nodes above the true MRCA, while fewer were found at descendant nodes below it. The remaining 8.7% were located on a different branch.

**Figure 2.**
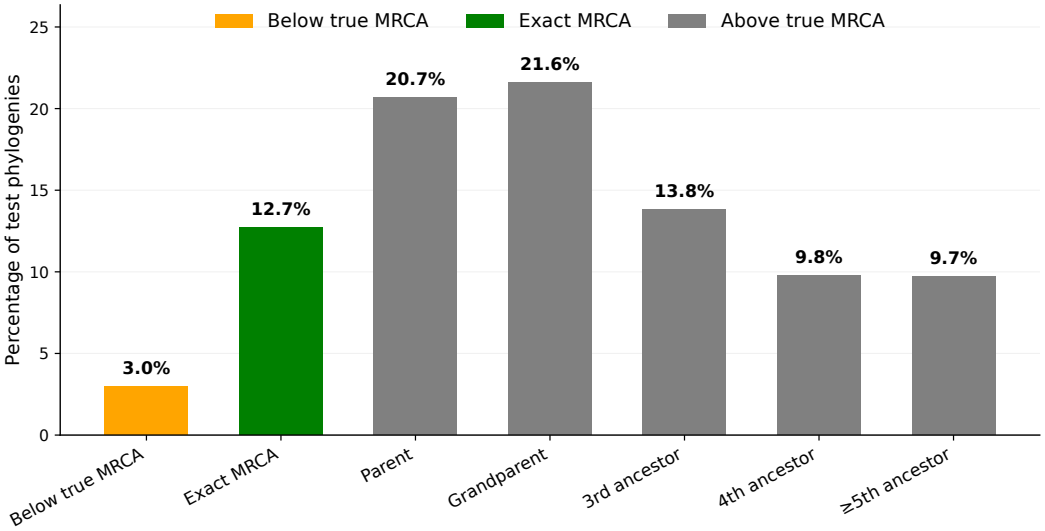
Distribution of predicted MRCA positions relative to the true MRCA across the independent test dataset. Positions above and below the true MRCA indicate ancestral and descendant predictions, respectively.

### Application to Empirical Data

In order to examine how well the proposed framework performs when used with data that is not from simulated datasets, we applied the trained model to a rooted SARS-CoV-2 phylogeny obtained from Santa Cruz County, California. Localized missingness was created by removing sequences from an internal clade, ensuring the relationships among the remaining sequences remained unchanged. We applied the recursive direction model, beam search, representative path selection, and MRCA estimation strategy to the observed phylogeny without retraining or changing any parameters. Table 5 shows the case study’s characteristics, and Figure 3 illustrates the predicted localization on the real phylogeny.

**Table 5.** Summary of the Santa Cruz SARS-CoV-2 case study.

| **Property** | **Value** |
|---|---|
| Full tree sequences | 72 |
| Observed tree sequences | 67 |
| Deleted sequences | 5 |
| Missing fraction within target clade | 21.7% |

**Figure 3.**
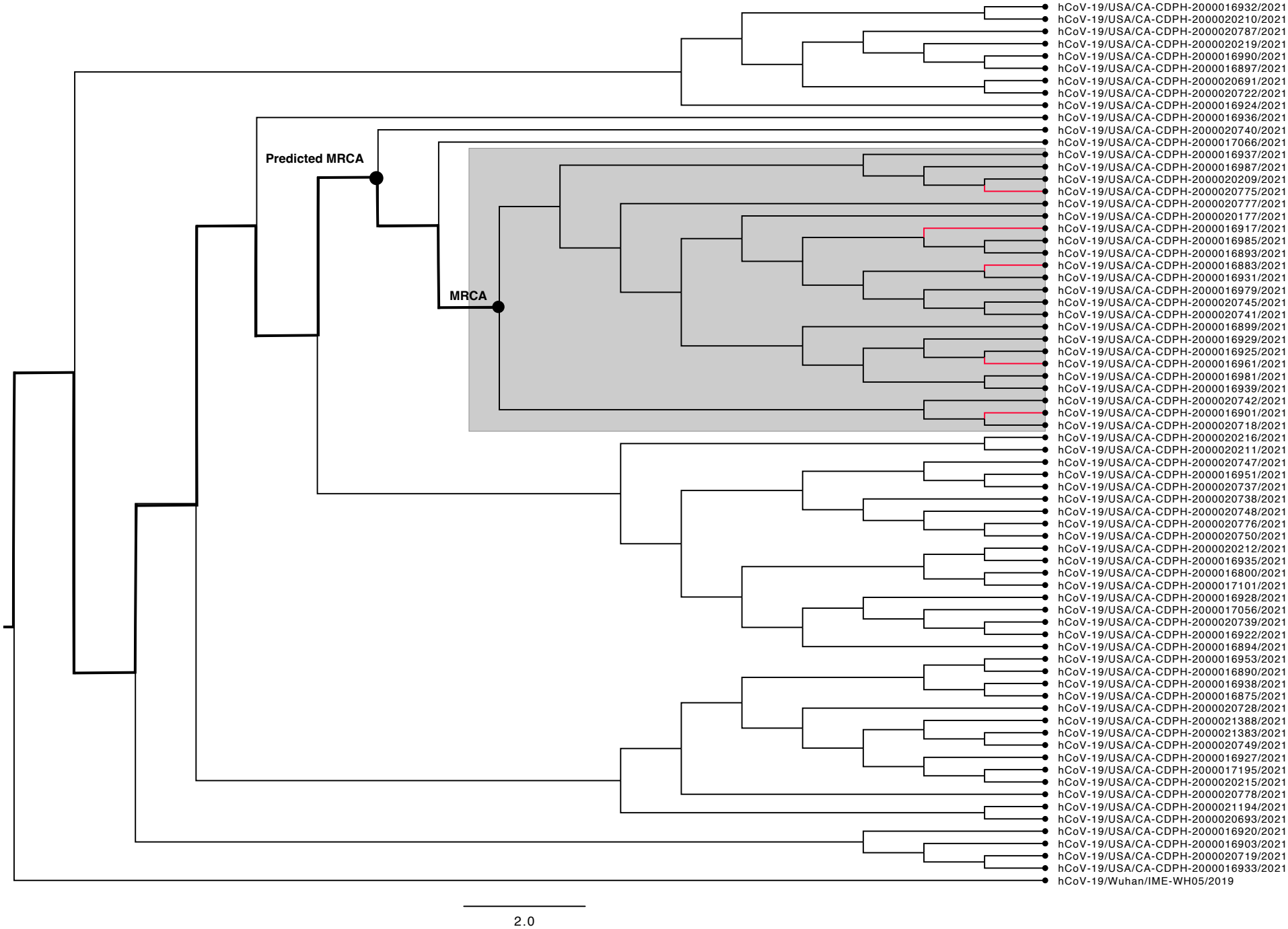
Application of the proposed framework to a rooted SARS-CoV-2 phylogeny from Santa Cruz County. The gray shaded region marks the simulated missing clade. Red terminal branches indicate the sequences removed from the complete phylogeny to simulate localized missingness. Bold branches show the predicted localization path from the root to the predicted MRCA. The true MRCA of the missing clade is shown for comparison.

The predicted localization path accurately found the ancestral lineage that led to the simulated missing clade. This result shows that the proposed framework can locate missing regions introduced into an empirical pathogen phylogeny.

## Discussion

Our results indicate that gaps resulting from localized missingness can produce a detectable structural signal within an observed phylogeny. Even in the absence of the missing sequences, the remaining phylogenetic tree often contains sufficient information to restrict their probable origin to a limited set of ancestral locations. Analysis of the Most Recent Common Ancestor (MRCA) further demonstrates that the predicted ancestral location typically resides on the same evolutionary lineage as the actual MRCA. These findings support the use of MRCA estimation to position the predicted missing region within the appropriate lineage of the tree. In practical terms, this approach may assist public health teams in identifying lineages that are potentially underrepresented in genomic surveillance and enhance near-real-time understanding of the distribution and spread of pathogen variants within specific jurisdictions.

The recursive direction model achieved about 79% accuracy on unseen trees, despite evaluating multiple learning algorithms and feature formulations. This suggests that some branching decisions are difficult to resolve from the structure of the observed phylogeny alone. When descendants are removed, the two child subtrees at an internal node often retain similar topological and branch-length characteristics, providing limited information about which branch leads toward the missing region. Maintaining multiple candidate trajectories addresses this uncertainty and ensures that the affected lineage remains among the reported candidates, even when individual direction predictions are uncertain or incorrect. Further improvements in localization performance will likely depend on improving robustness at branching points where the observed phylogeny provides weak or ambiguous structural evidence.

The present formulation deals with partial and local missingness, which occurs when only a subset of the descendants from an ancestral lineage are missing while the other descendants appear in the observed phylogeny. The part of the lineage that is retained provides information about structure that helps to locate the missing section. It has not yet been determined whether this signal remains informative in more complicated patterns of missingness, such as those affecting multiple lineages. Assessing more complex patterns alongside larger, more diverse phylogenies is essential to determine when the proposed approach remains reliable.

In practice, the framework could help prioritize regions of a pathogen phylogeny likely associated with unobserved evolutionary diversity, which may be linked to clinically meaningful differences, as seen among some COVID-19 variants of concern. By examining these candidate regions together with the sample metadata, it might be possible to detect possible gaps in terms of geographic, temporal, or population coverage. These findings can also be compared with clinical reports of unexplained changes in how severe a disease becomes. Using both types of information can help guide further sequencing or review of current surveillance data, especially in areas where missing diversity is likely. Identifying these missing regions can show where incomplete sampling might affect how we interpret later evolutionary or transmission studies.

## Data Availability

The SARS-CoV-2 sequences used to construct the 35 California county phylogenies were obtained from GISAID EpiCoV (https://gisaid.org). The GISAID EPI SET identifiers and corresponding DOIs for each county, along with the Wuhan sequence used for rooting, are listed in Supplementary Table S2. Sequence accession numbers and contributor acknowledgments are available through these GISAID records.

## Code Availability

The source code used to implement the methods and reproduce the analyses presented in this study is available at https://github.com/TaraKhodaei/phylo-missingness-localization.

## Author Contributions Statement

M.K. developed the computational framework, implemented the algorithms, performed the simulations and analyses, prepared the figures, and drafted the manuscript. N.H. co-supervised the research, provided feedback on its public-health relevance, and reviewed the manuscript. A.R. supervised the research, contributed to the study design and methodology, and revised the manuscript. All authors reviewed and approved the final manuscript.

## Acknowledgments

We gratefully acknowledge the authors and the originating and submitting laboratories that contributed the SARS-CoV-2 sequence data and associated metadata made available through GISAID EpiCoV and used in this study. Generative AI was used to improve the language and clarity of the manuscript. The authors reviewed and verified the final text.

