## Supplementary Material for "Localizing Unsampled Regions in Partially Sampled Pathogen Phylogenies Using Machine Learning"

Table S1. Definitions of the 52 structural features used by the recursive direction classifier. At each decision node, one child branch is evaluated as the candidate and the other as its sibling; the subtrees rooted at these children are referred to as the candidate and sibling subtrees.

| Feature | Definition |
|---|---|
| Observed-tree tip count | Total number of tips in the observed phylogeny. |
| Decision-node subtree tip count | Number of observed tips descending from the current decision node. |
| Candidate-subtree tip count | Number of observed tips descending from the candidate child. |
| Candidate-subtree height | Maximum branch-length distance from the candidate child to any tip in its subtree. |
| Candidate-subtree total branch length | Sum of all branch lengths within the candidate subtree. |
| Candidate-subtree mean branch length | Mean branch length across branches within the candidate subtree. |
| Candidate-subtree branch-length variance | Variance of branch lengths within the candidate subtree. |
| Candidate-subtree mean terminal branch length | Mean length of branches terminating at tips within the candidate subtree. |
| Candidate-subtree maximum terminal branch length | Maximum length among branches terminating at tips within the candidate subtree. |
| Total branch length per candidate tip | Candidate-subtree total branch length divided by candidate-subtree tip count. |
| Candidate-subtree height per tip | Candidate-subtree height divided by candidate-subtree tip count. |
| Candidate tip density by subtree height | Candidate-subtree tip count divided by candidate-subtree height. |
| Candidate tip density by total branch length | Candidate-subtree tip count divided by candidate-subtree total branch length. |
| Candidate fraction of decision-node tips | Proportion of tips below the decision node that are contained in the candidate subtree. |
| Candidate height relative to decision-node height | Candidate-subtree height divided by the height of the decision-node subtree. |
| Candidate branch length relative to decision-node branch length | Candidate-subtree total branch length divided by the total branch length of the decision-node subtree. |
| Candidate fraction of observed-tree tips | Proportion of all observed-tree tips contained in the candidate subtree. |
| Candidate height relative to observed-tree height | Candidate-subtree height divided by the height of the observed tree. |
| Candidate branch length relative to observed-tree branch length | Candidate-subtree total branch length divided by the total branch length of the observed tree. |
| Decision-node topological depth | Number of edges from the observed-tree root to the decision node. |
| Candidate-child topological depth | Number of edges from the observed-tree root to the candidate child. |
| Root-to-decision-node branch-length distance | Sum of branch lengths along the path from the observed-tree root to the decision node. |
| Root-to-candidate-child branch-length distance | Sum of branch lengths along the path from the observed-tree root to the candidate child. |
| Candidate smaller than sibling by tip count | Binary indicator equal to 1 when the candidate subtree contains fewer tips than the sibling subtree, and 0 otherwise. |
| Candidate smaller than sibling by total branch length | Binary indicator equal to 1 when the candidate subtree has smaller total branch length than the sibling subtree, and 0 otherwise. |
| Candidate share of sibling-pair tips | Candidate-subtree tip count as a proportion of the combined tip count of the candidate and sibling subtrees. |
| Candidate share of sibling-pair total branch length | Candidate-subtree total branch length as a proportion of the combined total branch length of the candidate and sibling subtrees. |
| Sibling-pair tip-count imbalance | Absolute difference in tip count between the candidate and sibling subtrees, normalized by their combined tip count. |
| Sibling-pair total-branch-length imbalance | Absolute difference in total branch length between the candidate and sibling subtrees, normalized by their combined total branch length. |
| Candidate terminal-branch fraction | Proportion of candidate-subtree total branch length contributed by terminal branches. |
| Sibling terminal-branch fraction | Proportion of sibling-subtree total branch length contributed by terminal branches. |

**Table S1.** Continued.

| Feature | Definition |
|---|---|
| Candidate internal tip-count balance | Ratio of the smaller to the larger tip count of the candidate's two immediate child subtrees. |
| Candidate internal height balance | Ratio of the smaller to the larger subtree height of the candidate's two immediate child subtrees. |
| Candidate internal total-branch-length balance | Ratio of the smaller to the larger total branch length of the candidate's two immediate child subtrees. |
| Candidate–sibling tip-count difference | Candidate-subtree tip count minus sibling-subtree tip count. |
| Candidate–sibling subtree-height difference | Candidate-subtree height minus sibling-subtree height. |
| Candidate–sibling total-branch-length difference | Candidate-subtree total branch length minus sibling-subtree total branch length. |
| Candidate–sibling mean terminal-branch-length difference | Candidate-subtree mean terminal branch length minus sibling-subtree mean terminal branch length. |
| Candidate–sibling maximum terminal-branch-length difference | Candidate-subtree maximum terminal branch length minus sibling-subtree maximum terminal branch length. |
| Candidate–sibling total-branch-length-per-tip difference | Candidate-subtree total branch length per tip minus the corresponding sibling-subtree value. |
| Candidate–sibling height-per-tip difference | Candidate-subtree height per tip minus the corresponding sibling-subtree value. |
| Candidate–sibling tip-density-by-height difference | Candidate tip density by subtree height minus the corresponding sibling-subtree value. |
| Candidate–sibling tip-density-by-branch-length difference | Candidate tip density by total branch length minus the corresponding sibling-subtree value. |
| Candidate-to-sibling tip-count ratio | Candidate-subtree tip count divided by sibling-subtree tip count. |
| Candidate-to-sibling subtree-height ratio | Candidate-subtree height divided by sibling-subtree height. |
| Candidate-to-sibling total-branch-length ratio | Candidate-subtree total branch length divided by sibling-subtree total branch length. |
| Candidate-to-sibling mean terminal-branch-length ratio | Candidate-subtree mean terminal branch length divided by sibling-subtree mean terminal branch length. |
| Candidate-to-sibling maximum terminal-branch-length ratio | Candidate-subtree maximum terminal branch length divided by sibling-subtree maximum terminal branch length. |
| Candidate-to-sibling total-branch-length-per-tip ratio | Candidate-subtree total branch length per tip divided by the corresponding sibling-subtree value. |
| Candidate-to-sibling height-per-tip ratio | Candidate-subtree height per tip divided by the corresponding sibling-subtree value. |
| Candidate-to-sibling tip-density-by-height ratio | Candidate tip density by subtree height divided by the corresponding sibling-subtree value. |
| Candidate-to-sibling tip-density-by-branch-length ratio | Candidate tip density by total branch length divided by the corresponding sibling-subtree value. |

Table S2. GISAID EPI_SET identifiers and persistent DOIs for SARS-CoV-2 sequences used in the 35 California county phylogenies. The Wuhan sequence used as an outgroup for rooting is listed separately.

| County / dataset | Sequences | Collection dates | GISAID EPI_SET | DOI |
|---|---|---|---|---|
| Butte County | 13 | Jan 2–6, 2021 | EPI_SET_261020fh | 10.55876/gis8.261020fh |
| Calaveras County | 13 | Jan 3–7, 2021 | EPI_SET_261020st | 10.55876/gis8.261020st |
| Contra Costa County | 211 | Jan 1–7, 2021 | EPI_SET_261020ow | 10.55876/gis8.261020ow |
| Fresno County | 162 | Jan 2–6, 2021 | EPI_SET_261020ut | 10.55876/gis8.261020ut |
| Humboldt County | 37 | Jan 3–7, 2021 | EPI_SET_261020pb | 10.55876/gis8.261020pb |
| Imperial County | 62 | Jan 4–7, 2021 | EPI_SET_261020hc | 10.55876/gis8.261020hc |
| Kern County | 47 | Jan 2–7, 2021 | EPI_SET_261020em | 10.55876/gis8.261020em |
| Kings County | 62 | Jan 3–7, 2021 | EPI_SET_261020nc | 10.55876/gis8.261020nc |
| Madera County | 12 | Jan 2–5, 2021 | EPI_SET_261020og | 10.55876/gis8.261020og |
| Marin County | 17 | Jan 2–5, 2021 | EPI_SET_261020yf | 10.55876/gis8.261020yf |
| Mendocino County | 41 | Jan 2–7, 2021 | EPI_SET_261020fy | 10.55876/gis8.261020fy |
| Merced County | 57 | Jan 3–6, 2021 | EPI_SET_261020th | 10.55876/gis8.261020th |
| Mono County | 19 | Jan 7, 2021 | EPI_SET_261020fv | 10.55876/gis8.261020fv |
| Monterey County | 202 | Jan 1–7, 2021 | EPI_SET_261020rw | 10.55876/gis8.261020rw |
| Nevada County | 12 | Jan 4–5, 2021 | EPI_SET_261020bf | 10.55876/gis8.261020bf |
| Orange County | 139 | Jan 1–7, 2021 | EPI_SET_261020ew | 10.55876/gis8.261020ew |
| Placer County | 22 | Jan 1–5, 2021 | EPI_SET_261020qg | 10.55876/gis8.261020qg |
| San Bernardino County | 183 | Jan 2–7, 2021 | EPI_SET_261020ef | 10.55876/gis8.261020ef |
| San Diego County | 35 | Jan 4–7, 2021 | EPI_SET_261020qo | 10.55876/gis8.261020qo |
| San Francisco County | 19 | Jan 2–7, 2021 | EPI_SET_261020hr | 10.55876/gis8.261020hr |
| San Joaquin County | 56 | Jan 2–6, 2021 | EPI_SET_261020vx | 10.55876/gis8.261020vx |
| San Luis Obispo County | 86 | Jan 3–7, 2021 | EPI_SET_261020sk | 10.55876/gis8.261020sk |
| San Mateo County | 18 | Jan 2–7, 2021 | EPI_SET_261020yr | 10.55876/gis8.261020yr |
| Santa Barbara County | 148 | Jan 3–6, 2021 | EPI_SET_261020af | 10.55876/gis8.261020af |
| Santa Clara County | 126 | Jan 1–7, 2021 | EPI_SET_261020te | 10.55876/gis8.261020te |
| Santa Cruz County | 71 | Jan 2–6, 2021 | EPI_SET_261020da | 10.55876/gis8.261020da |
| Shasta County | 12 | Jan 2–6, 2021 | EPI_SET_261020of | 10.55876/gis8.261020of |
| Solano County | 32 | Jan 2–6, 2021 | EPI_SET_261020bw | 10.55876/gis8.261020bw |
| Sonoma County | 42 | Jan 1–7, 2021 | EPI_SET_261020my | 10.55876/gis8.261020my |
| Stanislaus County | 93 | Jan 2–6, 2021 | EPI_SET_261020pq | 10.55876/gis8.261020pq |
| Sutter County | 18 | Jan 2–7, 2021 | EPI_SET_261020zg | 10.55876/gis8.261020zg |
| Tehama County | 20 | Jan 4–7, 2021 | EPI_SET_261020uv | 10.55876/gis8.261020uv |
| Tulare County | 83 | Jan 1–5, 2021 | EPI_SET_261020rt | 10.55876/gis8.261020rt |
| Tuolumne County | 15 | Jan 3–6, 2021 | EPI_SET_261020np | 10.55876/gis8.261020np |
| Ventura County | 12 | Jan 4–6, 2021 | EPI_SET_261020dx | 10.55876/gis8.261020dx |
| Wuhan outgroup | 1 | Dec 30, 2019 | EPI_SET_261020cs | 10.55876/gis8.261020cs |

*Note.* The 35 county datasets contain 2,197 sequences. An additional Wuhan sequence (hCoV-19/Wuhan/IME-WH05/2019; EPI_ISL_529217) was used as an outgroup for rooting, giving 2,198 sequences in total. Each DOI provides access to the corresponding GISAID dataset and contributor acknowledgments.
